# TRIB2 supports a high glycolytic phenotype in melanoma cells

**DOI:** 10.1101/2025.11.18.689112

**Authors:** Victor Mayoral-Varo, Ana Luísa De Sousa-Coelho, Carla Lopes, Óscar H. Martínez-Costa, Paulo J. Oliveira, Juan J. Aragón, Bibiana I. Ferreira, Wolfgang Link

**Affiliations:** Sols-Morreale Biomedical Research Institute (IIBM), Spanish National Research Council (CSIC), Universidad Autónoma de Madrid (UAM), Madrid, Spain; Faculty of Health Sciences - HM Hospitals, Camilo Jose Cela University. Villanueva de la Cañada 28692, Madrid, España; HM Hospitals Health Research Institute, Madrid 28015, Spain; Algarve Biomedical Center Research Institute (ABC-Ri), Campus de Gambelas, 8005-139 Faro, Portugal; Escola Superior de Saúde (ESS), Universidade do Algarve, Campus de Gambelas, 8005-139 Faro, Portugal; Center for Neuroscience and Cell Biology (CNC), Centre for Innovative Biomedicine and Biotechnology (CIBB), University of Coimbra, Coimbra, Portugal; Department of Biochemistry, Faculty of Medicine, Universidad Autónoma de Madrid (UAM), Arzobispo Morcillo, 4, 28029 Madrid, Spain; Faculdade de Medicina e Ciências Biomédicas (FMCB), Universidade do Algarve, Campus de Gambelas, 8005-139 Faro, Portugal

**Keywords:** Cancer metabolism, therapy resistance, TRIB2, Tribbles

## Abstract

Metabolic adaptation plays a crucial role in driving the progression of human melanoma contributing to therapy resistance and disease relapse. The pseudo serine/threonine kinase Tribbles homolog 2 (TRIB2) is prominently expressed in melanoma tissue, promoting resistance to anti-cancer treatments and correlating with unfavorable clinical outcomes. Despite this understanding, the impact of TRIB2 expression on melanoma’s metabolic profile remains unexplored. Here we use UACC-62 melanoma cells which exhibit substantial endogenous expression of TRIB2 as well as engineered isogenic TRIB2 knock out (KO) cells to assess the effect of the loss of TRIB2 on metabolism. Our findings reveal that TRIB2-KO cells display reduced glycolytic activity and heightened vulnerability to pharmacological inhibition of oxidative metabolism, contrasting with the parental cell line. These metabolic and phenotypic changes are driven by TRIB2’s coordinated regulation of multiple glycolytic genes, whose combined effect might produce the glycolytic phenotype observed in parental cells. Collectively, our results underscore important role of TRIB2 as a modulator of the metabolic shift implicated in therapy resistance. These insights highlight TRIB2 as a potential target for therapeutic intervention, aiming to counteract the metabolic adaptations driving melanoma’s resistance mechanisms.

**Highlights:**

1. Main problem in melanoma is the limited therapy efficacy due to resistance;
2. TRIB2 is a novel unexplored and druggable target;
3. TRIB2 expression contributes to increased glycolysis in melanoma cells;
4. TRIB2 is a driver of a metabolic switch that fuels therapy resistance.

## Introduction

Dysregulation of cellular metabolism is a hallmark of cancer [1]. Metabolic reprogramming may serve as a crucial mechanism of melanoma cells to survive treatment, with therapy resistance emerging as the principal cause behind treatment failure or relapse [2]. Melanoma, the most lethal type of skin cancer, has witnessed remarkable advances in therapy, notably mitogen-activated protein kinases (MAPK) and immune checkpoint inhibitors. However, the potential of these approaches is curtailed by drug resistance [3,4], accentuating the need for innovative melanoma treatments. Hence, the pursuit of novel anti-cancer agents or synergistic combinations, along with understanding key contributors becomes imperative. Furthermore, harnessing cellular metabolic vulnerabilities, once uncovered, offers promising avenues for novel therapeutic interventions.

Together with TRIB1 and TRIB3, Tribbles homolog 2 (TRIB2) belongs to the well-conserved mammalian Tribbles family of pseudokinases proteins [5]. There is evidence that TRIB2 overexpressing tumors show a malignant/chemoresistant phenotype in several types of cancer, spanning acute myeloid leukemia [6], chronic myelogenous leukemia [7], small cell lung cancer [8], glioblastoma (GBM) [9], colorectal cancer (CRC) [10,11], laryngeal squamous cell carcinoma [12], pancreatic cancer [13], and prostate cancer [14], suggesting TRIB2 as a therapeutic target for resistant tumors. In melanoma, we have identified TRIB2 as an oncogene and as a repressor of the metabolic master regulators forkhead box class O (FOXO) transcription factors through the activation of AKT [15]. Moreover, TRIB2 depletion in melanoma cells disrupts AKT/FOXO signaling in response to diverse drugs [16]. Therefore, we propose TRIB2 as a valuable biomarker for melanoma diagnose, progression assessment, and prediction of clinical responses to cancer treatments [17]. The aim of this study was to shed light on the role of TRIB2 in cellular metabolism, particularly its association with therapy resistance in melanoma.

## Results

### TRIB2-dependent cell viability in response to energy disruption drugs

Metformin (Metf), an oral anti-diabetic drug classified as a biguanide, has garnered attention for its potential antineoplastic properties across diverse cancer types including melanoma [18]. The antiproliferative effects of biguanides potentially stem from the activation of the nutrient sensor AMPK and the inhibition of complex I of the electron transport chain in the mitochondria (oxidative phosphorylation, OXPHOS), causing bioenergetic stress in cancer cells, and rendering them dependent on glycolysis for ATP production [19]. High endogenous TRIB2 protein levels were formerly detected in melanoma cells [15]. We previously developed isogenic TRIB2 knock out (KO) cells in UACC-62 melanoma cell line [16].

In parental UACC-62 cells (referred to as WT or wild-type), the inhibition of oxidative metabolism by Metformin (Metf, 50 mM) or Phenformin (Phen), 1 mM) for 48 hours showed a significant decrease in cells viability (Figure 1A,B – dark grey bars), as previously shown [20]. Notably, we found that TRIB2-depleted cells were more susceptible to the treatment with both biguanides, as viability of TRIB2-KO cells was further decreased in response to Metf and Phen (Figure 1A, B – light colored bars). This nuanced response may suggest that TRIB2-expressing cells rely more on glycolytic pathways than TRIB2-KO cells, as WT cells were partially more resistant to biguanides treatment.

**Figure 1.**
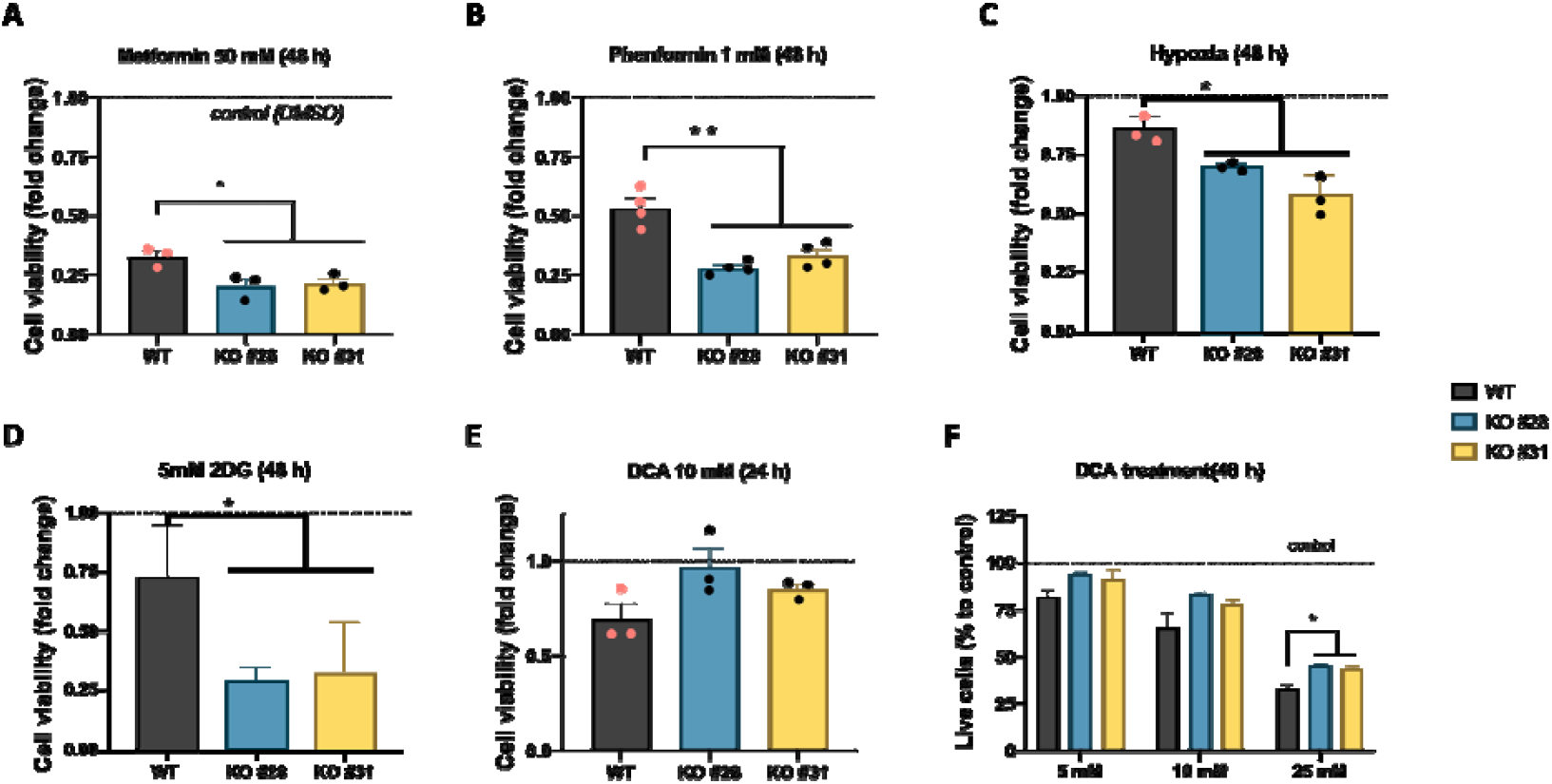
TRIB2-dependent cell viability in response to energy disruption drugs. (**A-E**) Cellular viability in WT and TRIB2-KO clones #28 and #31, obtained from MTT colorimetric assays, in response to 50 mM Metformin (**A**), 1 mM Phenformin (**B**), Hypoxia 1% O_2_ (**C**), 5 mM 2-Deoxiglucose (**D**) for 48 h and 10 mM dichloroacetic acid (DCA) for 24 h (**E**) (n=4 independent experiments performed in triplicate), as fold change to control (cells treated with vehicle DMSO 0.1%). (**F**) Proportion of live cells based on the trypan blue exclusion cell death assay in WT and TRIB2-KO #28 and #31, as percentage to each control cells, after 48 h treatment with increasing concentration of DCA (5, 10 and 25 mM) (n=3 independent experiments performed in triplicate). Values are mean +-standard error mean (SEM). ANOVA with Tukey’s multiple comparisons test *p* values indicated as ^*^ (*p*<0.05) or ^**^ (*p*<0.01).

To further evaluate whether TRIB2 affects cellular tolerance to metabolic stress, we exposed cells to acute hypoxia (1% O_2_, 48 h). TRIB2-KO cells exhibited a significant reduction in MTT signal compared with WT cells (Figure 1C), suggesting impaired survival under low-oxygen conditions. Because MTT reflects metabolic reductive capacity in addition to cell number, this decrease likely indicates both diminished metabolic activity and reduced viability in KO cells. We next examined cell viability following glycolysis inhibition using the hexokinase inhibitor 2-deoxyglucose (2-DG) [21]. After 48 h of treatment, TRIB2-KO cells showed a significantly greater reduction in viability than WT cells (Figure 1D), consistent with their lower glycolytic capacity and reduced ability to compensate when glycolysis is acutely blocked.

Finally, when exposed to dichloroacetic acid (DCA), which selectively inhibits pyruvate dehydrogenase kinases (PDKs) and consequently impairs glycolysis favoring mitochondrial oxidative metabolism [22,23], KO cells displayed a modest decrease in viability, though not statistically significant (Figure 1E). A dose-dependent variance in cell death further underscored this difference, revealing enhanced survival of TRIB2-depleted cells upon glycolysis inhibition (Figure 1F). Our observations revealed that subjecting cells to a 48-hour treatment with 25 mM of DCA led to increased cell death in TRIB2-expressing (WT) cells, surpassing the response in TRIB2-KO cells (Figure 1F). These findings prompt our hypothesis that TRIB2 potentially plays a role in sustaining elevated glycolytic capacity in melanoma cells.

### Cells with high TRIB2 levels show preferential glycolytic metabolism

Given the influence of OXPHOS inhibition on cellular viability, contingent upon TRIB2 expression in UACC-62 cells, we decided to study whether TRIB2 was involved in their cellular metabolism. We found both increased glucose consumption and lactate production in WT cells compared to TRIB2-KO cells, cultured in high glucose media (Figure 2A, B). These results suggest that the depletion of TRIB2 in melanoma cells renders them less likely to use glucose as their preferential energetic substrate.

**Figure 2.**
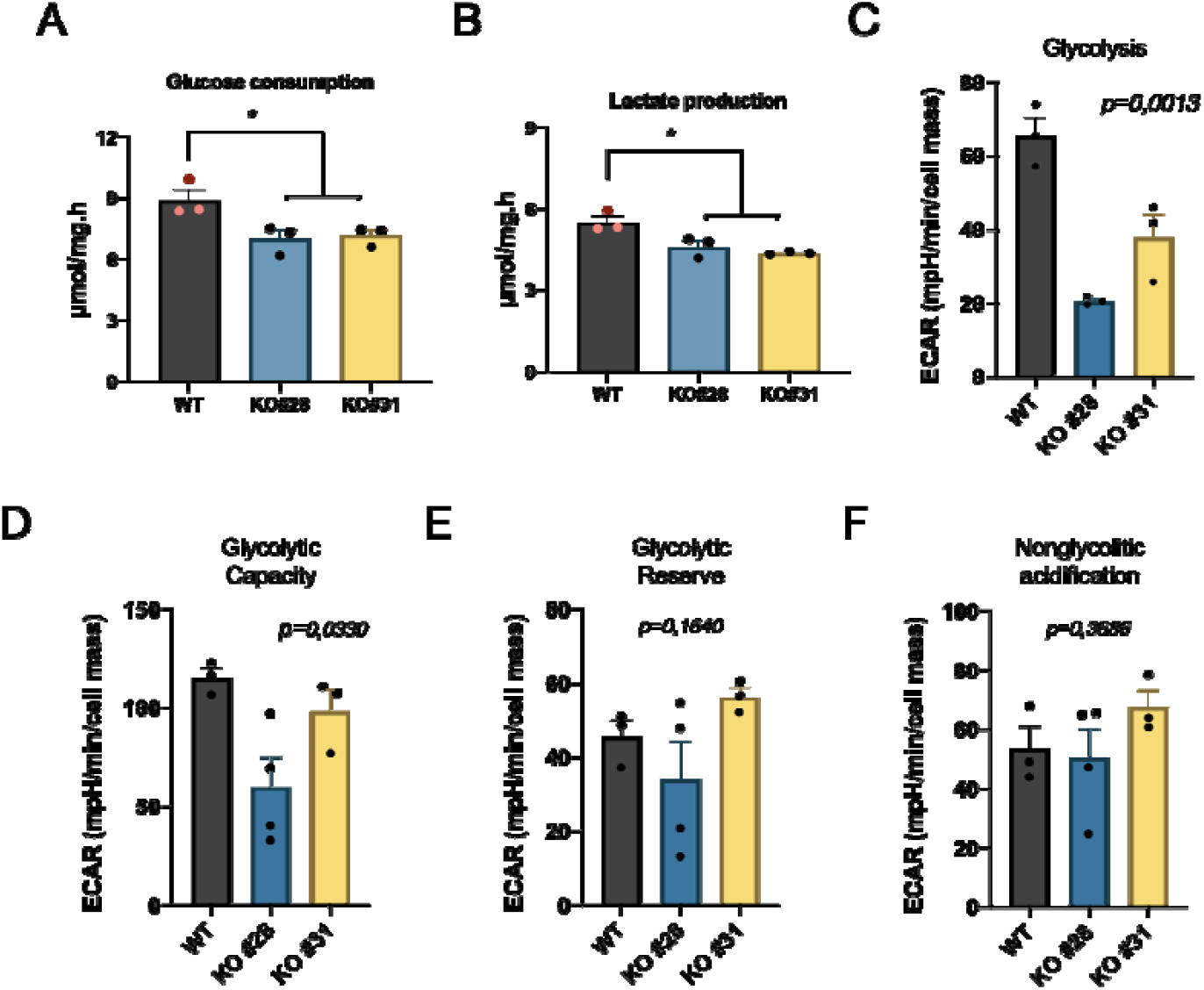
Cells with high TRIB2 levels show preferential glycolytic metabolism. (**A**) Glucose consumption and (**B**) lactate secretion, represented as µmol/mg.h, from parental (WT) and 2 independent clones of TRIB2-KO (#28 and #31) UACC-62 cells, cultured for 4 h in high glucose media after a 1-hour fasting period (DMEM without glucose), obtained from spectrometry-based enzymatic assays (n=3 independent assays performed in triplicate). (**C-F**) Individual parameters of glycolysis based on the extracellular acidification rate (ECAR) in WT and TRIB2-KO #28 and #31 cells, evaluated through the Seahorse bioanalyzer (n=3-4 assays performed with 5 replicates). Values are mean +-standard error mean (SEM). Tukey’s multiple comparisons test *p* values indicated as ^*^ (*p*<0.05) (**A**,**B**) or ANOVA *p* values shown (**C-F**).

To determine TRIB2-induced global changes in metabolic features, we evaluated the oxygen consumption rates (OCR), in parallel with extracellular acidification rate (ECAR), which allow the real-time analysis of both oxidative and glycolytic metabolism in live cells, respectively. Functional assays revealed decreased glycolysis and glycolytic capacity in TRIB2-KO cells, compared to WT cells (Figure 2C, D, Figure S1A). Notably, these differences did not extend to their glycolytic reserve or levels of non-glycolytic acidification, as statistical significance was absent (Figure 2E, F). Here, we recognized for the first time that cells with high TRIB2 levels have a glycolytic preference for survival.

### TRIB2 expression does not alter mitochondrial respiration

Based on the observation that cells without TRIB2 rely less on glycolysis, we anticipated KO cells would show increased oxidative metabolism and mitochondrial function, as a compensatory mechanism. However, we found no significant differences in any of the measurements performed based on the OCR, namely basal respiration, maximal respiratory capacity, ATP linked respiration, nor in spare respiratory capacity and proton leak, between WT and TRIB2-KO cells (Figure 3A-E; Figure S1B).

**Figure 3.**
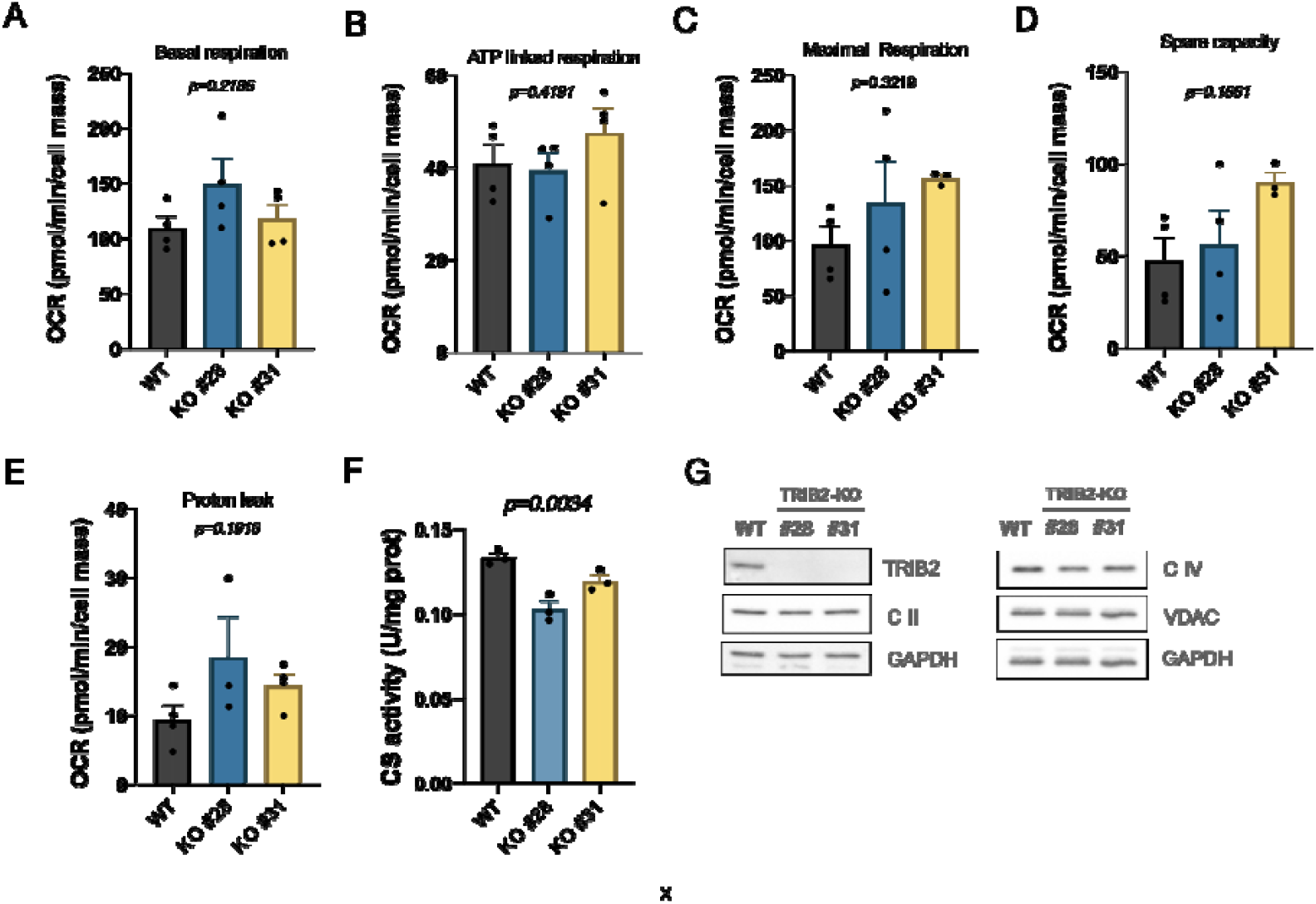
Mitochondrial function does not depend on TRIB2 expression. (**A-E**) Individual parameters of mitochondrial function based on the oxygen consumption rate (OCR) in WT and TRIB2-KO #28 and #31 cells, evaluated through the Seahorse bioanalyzer (n=3-4 assays performed with 5 replicates). (**F**) Citrate synthase activity in WT and TRIB2-KO #28 and #31 cells (n=3 assays performed in triplicate). Values are mean +-standard error mean (SEM). ANOVA test *p* values are indicated. (**G**) Mitochondrial proteins levels in whole cell extracts from parental (WT) and TRIB2-KO (clone #28 and #31) UACC-62 melanoma cell line, using GAPDH as loading control.

As a surrogate of mitochondrial mass and function, we also assessed the citrate synthase (CS) activity and mitochondrial protein levels. While CS activity was decreased in TRIB2-KO cells, compared to WT cells (Figure 3F), the protein levels of OXPHOS complex II (UQCRC2) and complex IV (COX II), and Voltage Dependent Anion Channel 1 (VDAC1) were similar between genotypes (Figure 3G), suggesting other energetic sources are fueling the cell without changes in their mitochondrial activity.

### Altered expression of metabolic genes in TRIB2-KO cells

Several key enzymes and transporters orchestrate intricate glycolytic processes [24]. To identify the pathways and gene expression profiles potentially modulated by TRIB2, we selected and analyzed the transcript levels of different possible contributors aiming to decipher the underpinning of the observed distinct response (Figure 4).

**Figure 4.**
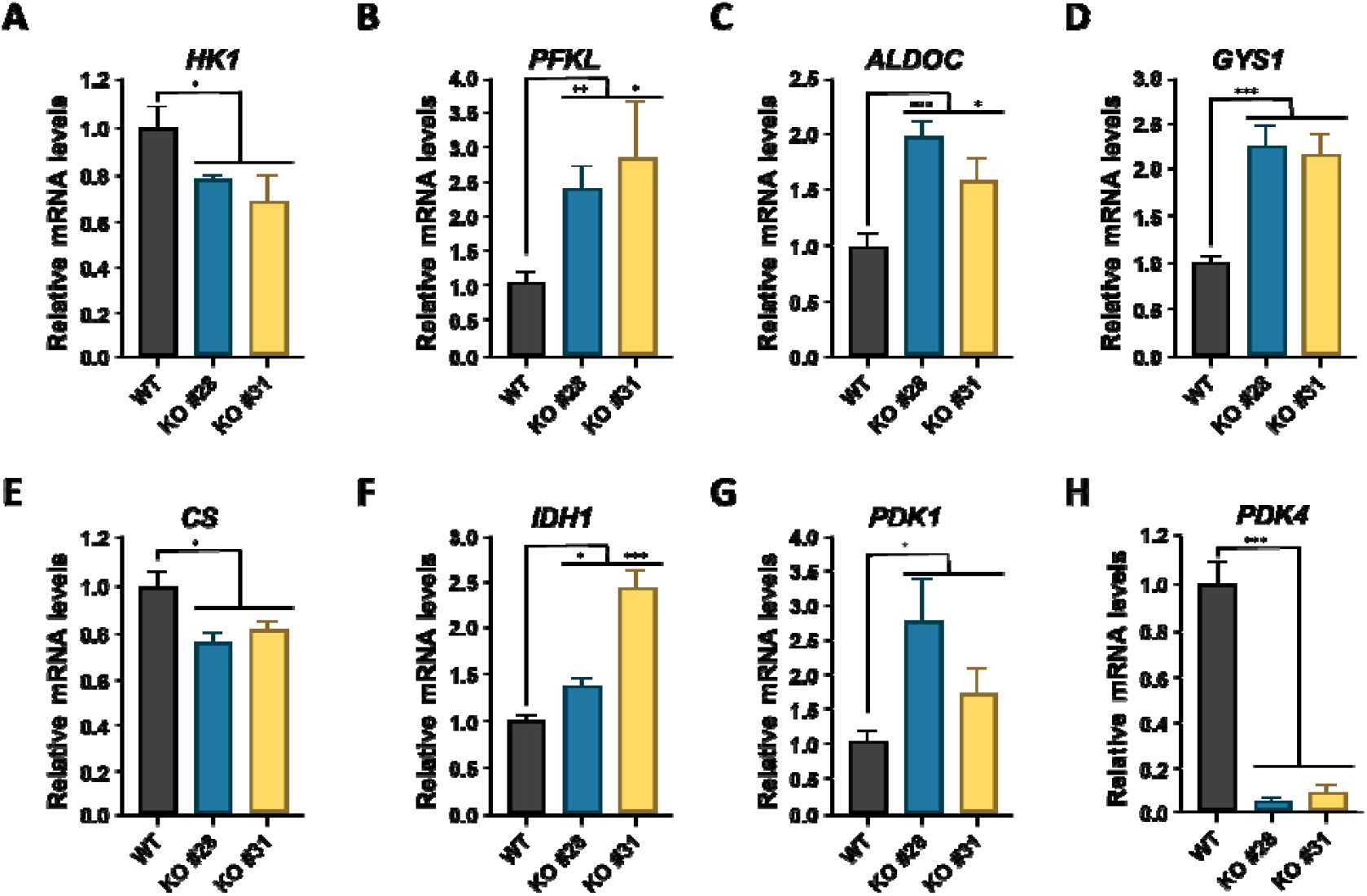
Depletion of TRIB2 alters the expression of genes involved in metabolism. (**A-H**) Relative mRNA expression of hexokinase 1 (HK1), phosphofructokinase liver type (PFKL), aldolase C (ALDOC), glycogen synthase 1 (GYS1), citrate synthase (CS), isocitrate dehydrogenase 1 (IDH1), pyruvate dehydrogenase kinase 1 (PDK1) and pyruvate dehydrogenase kinase 4 (PDK4),, normalized to GAPDH, in parental (WT) and 2 independent clones of TRIB2-KO (#28 and #31) UACC-62 cells (n=5-7 independent assays). Values are mean +-standard error mean (SEM) of fold change to WT. ANOVA was performed and *p* values are indicated in each graph.

Genes whose expression remained unchanged included GLUT1 (SLC2A1), PGM1, PKM, GPI, as well as regulators associated with mitochondrial oxidative metabolism (PGC1A and CPT1A) and the glycogen regulatory kinase GSK3B (Figure S2). These results indicate that TRIB2 deletion does not broadly alter basal expression of these metabolic components.

In contrast, several glycolytic and biosynthetic genes exhibited clear differences between WT and TRIB2-KO cells. Hexokinase 1 (HK1) mRNA levels were significantly reduced in both KO clones compared with WT (Figure 4A), consistent with a decrease in the first committed step of glycolysis. Conversely, phosphofructokinase liver type (PFKL) expression was elevated in KO cells (Figure 4B), suggesting an increase in upper-glycolytic flux downstream of hexokinase. Two additional glycolytic or branching-pathway genes also showed consistent upregulation: ALDOC, encoding the aldolase C enzyme (Figure 4C), and GYS1, encoding glycogen synthase 1 (Figure 4D). To determine whether TRIB2 deletion also affected mitochondrial metabolism, we examined two enzymes of the tricarboxylic acid (TCA) cycle: citrate synthase (CS), which catalyzes the first committed step of the cycle, and isocitrate dehydrogenase 1 (IDH1). CS expression was reduced in TRIB2-KO cells, while IDH1 levels were increased (Figure 4E–F), indicating that TRIB2 influences not only glycolytic regulation but also enzymes associated with mitochondrial carbon flux.

We next examined the pyruvate dehydrogenase kinase (PDK) family, which controls activity of the pyruvate dehydrogenase complex (PDC), a key regulatory node linking glycolysis to the tricarboxylic acid cycle [25]. PDK1 mRNA was moderately increased in both TRIB2-KO clones (Figure 4G), whereas PDK4 expression was markedly reduced in KO cells compared with WT (Figure 4H). These reciprocal changes in PDK isoforms suggest a shift in the regulatory balance controlling pyruvate entry into the mitochondria.

Finally, we assessed whether modulation of PDK isoforms affected PDC activity by measuring phosphorylation of the PDH E1α subunit at serine 300. No differences in P-PDH (S300) levels were detected between WT and KO cells (Figure S3A), indicating that PDC phosphorylation may be buffered by compensatory mechanisms despite the strong downregulation of PDK4. In agreement with this, protein levels of LDHA and PKM2 remained unchanged between genotypes (Figure S3B).

## Discussion

TRIB2, recognized as an oncogene which interacts with proteins involved in cell survival and proliferation [27], has previously demonstrated a distinct role in fueling the growth and survival of melanoma cells [15]. In line with this, our study revealed an enhanced glycolytic propensity in TRIB2-expressing cells compared to TRIB2-KO cells. This glycolytic bias was evident through the partial resistance of WT cells to biguanide treatment, which triggers a transition from oxidative phosphorylation to glycolysis. In-depth exploration of UACC-62 human melanoma cells bearing high endogenous TRIB2 expression unveiled increased glucose consumption and lactate production compared to KO cells. Notably, this shift was independent of LDHA, the enzyme that metabolizes pyruvate to produce lactate, which remained consistent between WT and KO cells.

While a majority of cancer cells are conventionally associated with increased glycolytic activity [28], the prospect of inhibiting glycolysis emerges as an appealing therapeutic strategy for specific cancer types or stages. Furthermore, resistance to therapies can usher in metabolic adaptations, offering an avenue for combinatorial treatments. This concept is exemplified by recent findings showing that mTOR-induced cancer drug resistance was linked to hypersensitivity to glycolytic inhibitors 2-deoxy-glucose (2DG) and dichloroacetate (DCA) [29], underscoring these exploitable metabolic vulnerabilities.

Beyond these global phenotypic differences, our data indicate that TRIB2 modulates melanoma metabolism through the coordinated regulation of multiple metabolic genes across several nodes of central carbon metabolism, rather than acting through a single dominant effector. TRIB2-KO cells exhibited reduced HK1 expression, but also increased levels of PFKL, ALDOC, and GYS1, suggesting that TRIB2 influences glycolytic flux through distributed transcriptional adjustments that collectively reduce the ability of KO cells to engage glycolysis.

Importantly, this regulatory influence extended into the tricarboxylic acid (TCA) cycle, where TRIB2 depletion resulted in reduced expression of citrate synthase (CS) and increased levels of IDH1. These enzymes represent key entry and progression points within the TCA cycle, and their altered expression provides a mechanistic explanation for the reduced glycolytic capacity and diminished lactate production observed in TRIB2-KO cells.

The existence of such a distributed metabolic signature aligns with recent evidence showing that metabolic rewiring in cancer is frequently driven by transcriptionally coordinated programs. For example, MYC simultaneously controls genes involved in glycolysis, mitochondrial biogenesis, and glutamine metabolism, establishing broad metabolic reprogramming [30]. Similarly, KRAS activates parallel anabolic pathways involving glycolysis, the pentose phosphate pathway, and TCA-cycle–linked anaplerosis [31]. PGC1α also illustrates how a single regulatory node can modulate dozens of mitochondrial and oxidative genes through a unified transcriptional network [32]. In light of these precedents, our findings support the notion that TRIB2 behaves as an upstream modulator of multiple metabolic routes in melanoma cells.

Within mammals, PDK isoforms exhibit tissue- or cell-type-specific patterns. Although PDK2, prevalent across various tissues, has been identified as exquisitely sensitive to DCA inhibition [33,34], our analysis of UACC-62 failed to detect endogenous levels of PDK2 by qPCR (not shown). In contrast, PDK4 intimately associated with physiological metabolic adaptability, garners high expression in organs such as the heart, skeletal muscle, liver, kidney, and pancreatic islets [35]. Intriguingly, PDK4’s role emerges as context-dependent, manifesting as either an oncogenic or tumor-suppressive factor depending on the tumor type, while also featuring implications in therapy resistance [36]. Although PDK4 was markedly reduced in TRIB2-KO cells, this observation alone does not explain the entirety of the metabolic phenotype. The concomitant increase of PDK1, the preserved phosphorylation of PDH, and the broader transcriptional modifications observed across glycolytic and TCA-cycle genes indicate that PDK4 downregulation is part of a more complex metabolic adjustment rather than the sole driver. Even with reduced levels of PDK4 in TRIB2-KO cells, the unmodified levels of P-PDH (S300), compared to WT cells, were not entirely unexpected. Tiersma *et al*. recently show that even after knocking down PDK isoforms individually, namely after downregulation of PDK4 in A375 melanoma cells, the phosphorylation levels of PDH remained comparable to control [37]. Consistently, mitochondrial function assessed by OCR remained unchanged across WT and TRIB2-KO cells, and no differences were found in the abundance of representative mitochondrial respiratory chain complexes. Together, these findings indicate that mitochondrial oxidative capacity remains intact and that the reduction in CS expression and activity reflects a specific metabolic adjustment rather than a global mitochondrial deficit.

The differential impact of DCA was indicated even at lower concentrations (5 and 10 mM), becoming statistically significant at 25 mM. Indeed, in other melanoma cell lines, the influence of DCA on cell proliferation after 48 hours was mainly noticeable only at in A375 cells either at 20 mM or above this concentration in Mewo cells [38]. Nevertheless, a 96-hour treatment led, to a minimum 25% reduction in melanoma cell viability with 10 mM of DCA [39]. Previous clinical studies revealed dose-limiting toxicity of that DCA, namely reversible peripheral neuropathy [40–43]. Based on our results, we predict that in tumors with high TRIB2 expression, lower concentrations and duration of DCA treatment can be used, which may contribute to reduce the adverse effects for patients, who will benefit from such therapeutic strategy.

Over the past three years, the literature has seen an influx of over 7,000 articles on glucose metabolism and cancer, with around 160 focused on melanoma. Notably, very few publications have delved into the relationship between Tribbles and glucose metabolism, with only one recently shedding light on TRIB2’s potential role in this process [44]. Liu *et al*. showed that in lung cancer cells, TRIB2 would interact with and regulate PKM2, the M2 isoform of pyruvate kinase, promoting the aerobic glycolysis [44]. In agreement with our results, glucose uptake and lactate production were increased in TRIB2-overexpressing cells [44]. However, while the authors showed that PKM2 levels were modulated by TRIB2 protein, in our experiments no differences in PKM2 levels were observed between in WT and UACC-62 cells, both at the mRNA and protein level. We also did not find differences in *GLUT1* transcript or LDHA protein levels, although these disparities could reflect the different type of tumor and should be further explored.

Altogether, our integrated transcriptional, metabolic, and functional analyses support a model in which TRIB2 promotes a glycolytic phenotype through distributed regulation of multiple metabolic genes that span glycolysis, the TCA cycle, and pyruvate utilization. Rather than controlling a single metabolic switch, TRIB2 appears to influence melanoma cell bioenergetics through a coordinated remodeling of several metabolic nodes, a mechanism analogous to those described for other oncogenic regulators such as MYC, KRAS, and PGC1α [30-32].

Given the greater glycolytic flux in TRIB2(+) cells, TRIB2 expression in melanoma and other tumors appears to instigate reliance on glycolysis. In this scenario, chemotherapy combined with biguanides would only benefit tumors with diminished TRIB2 levels. This insight is of great relevance, as the biguanide drug Phenformin is gaining renewed attention as a superior option to Metformin in melanoma [45], and is currently evaluated in clinical trials (NCT03026517). Conversely, co-inhibiting glycolysis (for instance through short-term treatment with DCA, which targets pyruvate dehydrogenase kinase, thereby promoting oxidative metabolism of pyruvate in the mitochondria [46,47]), would likely improve the response of melanoma patients with high levels of TRIB2.

In summary, our findings provide evidence for a potential role of TRIB2 in driving malignant cells towards an aerobic glycolytic phenotype, a hallmark of cancer, potentially contributing to the resistance to cancer therapies.

## Methods

### Cell lines and treatments

Isogenic melanoma cell lines, corresponding to WT [TRIB2(+)], and CRISPR/TRIB2-KO [TRIB2(-), undetectable TRIB2 protein, clones #28 and #31] UACC-62 cells were developed and described previously [16]. Briefly, after cells were transfected with two independent guide RNAs in plasmid vectors expressing Cas9 nuclease protein, single-cell clones were selected and expanded. Finally, the full deletion of the protein levels of TRIB2 were evaluated by Western blot.

Cells were cultured in standard conditions of temperature (37ºC) and CO_2_ (5%), and maintained in DMEM high glucose (25 mM glucose) (Cytiva SH30243.LS) supplemented with 10% fetal bovine serum (FBS) and antibiotics (A8943.0100, Panreac Applichem), as described previously [20]. As standard procedure, all cell lines were periodically tested for mycoplasma contamination.

Metformin (13118) and Phenformin (14997) were from Cayman Chemical Company. Sodium dichloroacetate (DCA) (347795) and 2-deoxyglucose (2-DG) (D8375) was purchased from Sigma-Aldrich. Metformin, DCA and 2-DG were dissolved in DMEM media, before being added to the cells in culture. Phenformin was dissolved in DMSO.

### Viability and cell death assays

Cells viability was determined using the 3-(4,5-dimethylthiazol-2-yl)-2,5-diphenyltetrazolium bromide (MTT) colorimetric assay or the cellular death trypan blue exclusion assay, as described previously [20].

### Protein extraction, quantification and Western blotting

After being washed with cold phosphate buffer saline (PBS), cells were lysed using CST buffer supplemented with phosphatase and protease inhibitors [20]. Proteins present at the lysates were quantified using a Bradford assay (J61522-K2 Alfa Aesar). Protein samples (30 µg) were mixed together with Laemmli Buffer and heated at 95□°C for 5□min, separated in 8% or 10% SDS/PAGE gels and electroblotted to polyvinylidene fluoride (PVDF) methanol-activated membrane (GE Healthcare Life Sciences). The membranes were blocked in Tris-buffered saline with 0.1% Tween® 20 detergent (TBS-T) supplemented with 5% non-fat milk, for at least 1Lh in room temperature, before labeled with primary antibodies diluted at the indicated concentrations in TBS-T 5% bovine serum albumin (BSA) overnight at 4□°C with gentle shaking. After washing in TBS-T buffer the membranes were incubated for at least 1□h in respective secondary antibodies at the appropriate dilutions. The protein bands were visualized using the iBright 1500 Imaging System (Invitrogen, Thermo Fisher Scientific).

The following primary antibodies were used: total PDHA1 (ab168379), VDAC (ab154856) and total OXPHOS human WB antibody cocktail (ab110411) from Abcam; Phospho PDH-E1 (pSer300) (AP1064) from Millipore, Calbiochem; and TRIB2 (D8P2X, #13533), LDHA (C4B5, #3582S) and PKM2 (D78A4, #4053S), from Cell Signaling Technology. Secondary antibodies were conjugated with horseradish peroxidase (GE Healthcare Life Sciences) and visualized by the ECL detection solution. Whenever possible based on the molecular size of the proteins, membranes were re-stained with different antibodies, including antibodies for protein loading control (rabbit anti-GAPDH, glyceraldehyde 3-phosphate dehydrogenase (sc-25778, Santa Cruz Biotechnology) and mouse anti-alpha Tubulin (T9026, Sigma-Aldrich). Secondary antibodies were ECL Mouse IgG, horseradish peroxidase (HRP)-linked whole Ab (from sheep) (NA931) and ECL Rabbit IgG, HRP-linked whole Ab (from donkey) (NA934).

### Lactate and glucose measurements

For glucose and lactate level measurements, melanoma cells were plated in 60 mm dishes at a final density of 5×10^5^ cells and incubated at 37ºC for 24 h. Cells were then placed in DMEM without glucose and glutamine for 1 h. After the fasting period, cells were placed back in complete DMEM (time 0) for additional 4 h (time 4). An aliquot of culture media was collected both at time 0 and time 4 and stored at −80°C until analysis. Cells were detached with PBS and stored at −80°C for protein extraction. Glucose and lactate levels present in the conditioned medium in culture were quantified using standard enzymatic techniques, subtracted to the initial levels (time 0), and normalized by protein amount (obtained and quantified as described above), as previously described [48].

### Citrate synthase activity assay

Citrate synthase (CS) activity was determined using 0.2 mM acetyl-CoA, 0.5 mM oxaloacetate (OAA), and 0.1 mM 5,5’-dithiobis(2-nitrobenzoic acid) DTNB in 100 mM Tris-HCl (pH 7.5) [49]. The reaction was followed spectrophotometrically at 412 nm at room temperature. Enzymatic activity was normalized by total protein amount.

### RNA extraction and gene expression analysis

For gene expression analysis, cells were washed with cold PBS, pelleted and stored at −20□°C until further processing. Cellular RNA was extracted using the E.Z.N.A. Total RNA Kit I, including DNase digestion (R6834-02, VWR). Total RNA concentration was estimated using a Nanodrop Spectrophotometer and 2 µg were reversely transcribed using the NZY First-Strand cDNA Synthesis Kit (MB12502, Nzytech) according to manufacturer instructions.

### Bioenergetics assay

The oxidative phosphorylation and glycolytic profile of UACC-62 melanoma cells were obtained by measuring the oxygen consumption rate (OCR) and extracellular acidification ratio (ECAR), on a Seahorse XF^e^96 Extracellular Flux Analyzer (Agilent Technologies, Germany). For OCR and ECAR, cells were seeded 12 000 cells/per well onto an XF96 Cell Culture Microplate and allowed to adhere for 24 h at 37ºC. A volume of 200 μL of calibration buffer was placed into each well of the sensor hydration microplate and the sensor cartridge placed onto the microplate. The plate was incubated with immersed sensors in a non-CO^2^ incubator overnight at 37°C. For OCR analysis, assay medium (DMEM D5030, Sigma) was supplemented with 4,5 g/L glucose, 4.0 mM L-glutamine and 1 mM sodium pyruvate, pH adjusted to 7.4, at 37°C; whilst for ECAR no glucose was added. The following day the cell medium was aspirated and gently each well was washed with assay medium at 37 °C. Then, 180 µL of assay medium at 37°C was added to each well and the plate placed into a non-CO^2^ incubator at 37°C for 1 h to reach equilibrium. To measure OCR, four compounds were added into successive injection ports, with the following final concentrations: A - 1 μM oligomycin; B - 1 μM carbonyl cyanide 4-(trifluoromethoxy)phenylhydrazone (FCCP); and C - 1 μM rotenone plus 1 μM antimycin A. For the ECAR analysis the following compounds were injected into the delivery ports: A - saturated concentration of glucose (10 mM), B - 1 μM oligomycin and C - 50 mM 2-deoxyglucose (2-DG). The protocol consists of three measurement cycles before the first injection and three measurement cycles after each port injection. Each measurement cycle consisted of 30 s mixing and 3 min of measurement. After the assay was completed, cells were fixed by adding 50 μL of 60 % trichloroacetic acid and stored for 24 h at 4ºC. Cell mass was quantified by Sulforhodamine B assay and used to normalize individual wells [50]. All the results analyses were performed using the Software Wave Desktop Version 2.2.

### Statistical analysis

Statistical analysis was performed using GraphPad Prism software (version 8.4.0, San Diego, CA, USA). ANOVA test was used to evaluate statistical differences between the WT and TRIB2-KO cells. When indicated, multiple comparisons were performed. *P*□<□0.05 was considered statistically significant.

## Acknowledgements

This work was supported by the Spanish Ministry of Science, Innovation and Universities through Grant PID2022-136654OB-I00 to WL, and Fundação para a Ciência e a Tecnologia (FCT) (PTDC/MED-ONC/4167/2020 “ENDURING”) to BIF.

## Author contributions

Ana Luísa De Sousa-Coelho: Conceptualization, Formal analysis, Investigation, Methodology, Project administration, Visualization, Writing - original draft, Writing - review & editing. Victor Mayoral-Varo: Formal analysis, Investigation, Methodology, Project administration, Funding acquisition, Writing - review & editing. Óscar H. Martínez-Costa: Investigation. Carla Lopes: Formal analysis, Investigation. Paulo J. Oliveira: Methodology, Resources. Juan J. Aragón: Methodology, Resources. Bibiana I. Ferreira: Funding acquisition, Resources, Writing - review & editing. Wolfgang Link: Funding acquisition, Project administration, Resources, Supervision, Writing - review & editing.

