## Supplementary material for "TRIB2 supports a high glycolytic phenotype in melanoma cells": Suppl. Material

### Supplementary figures

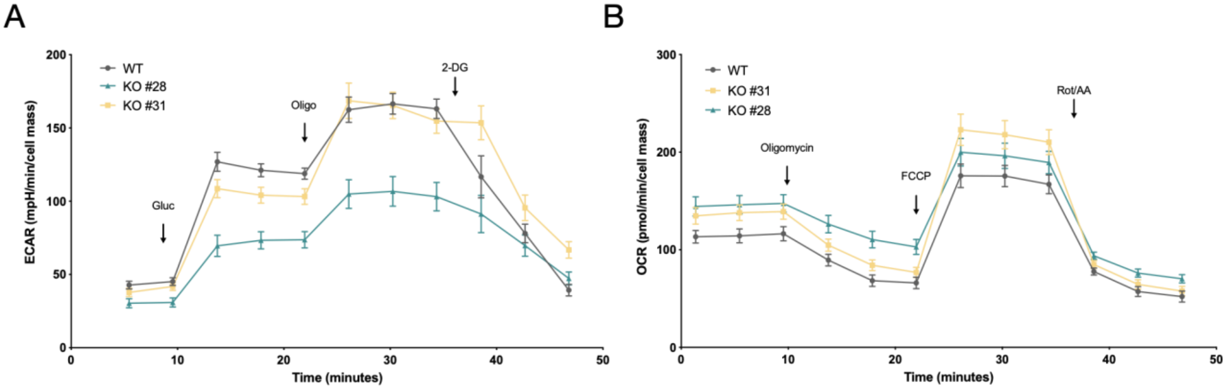


**Figure S1**. (**A**) Agilent Seahorse XF96 analysis of glycolytic function based on the extracellular acidification rate (ECAR) in WT and TRIB2-KO #28 and #31 cells, in basal media without glucose, and in response to glucose (Gluc), oligomycin (Oligo) and 2-deoxy-glucose (2-DG). (**B**) Agilent Seahorse XF96 analysis of mitochondrial respiration based on the oxygen consumption rate (OCR) in WT and TRIB2-KO #28 and #31 cells, in basal media and in response to oligomycin, carbonyl cyanide-4 (trifluoromethoxy) phenylhydrazone (FCCP) and a mix of rotenone (Rot) and antimycin A (AA). Values are mean +- standard error mean (SEM).

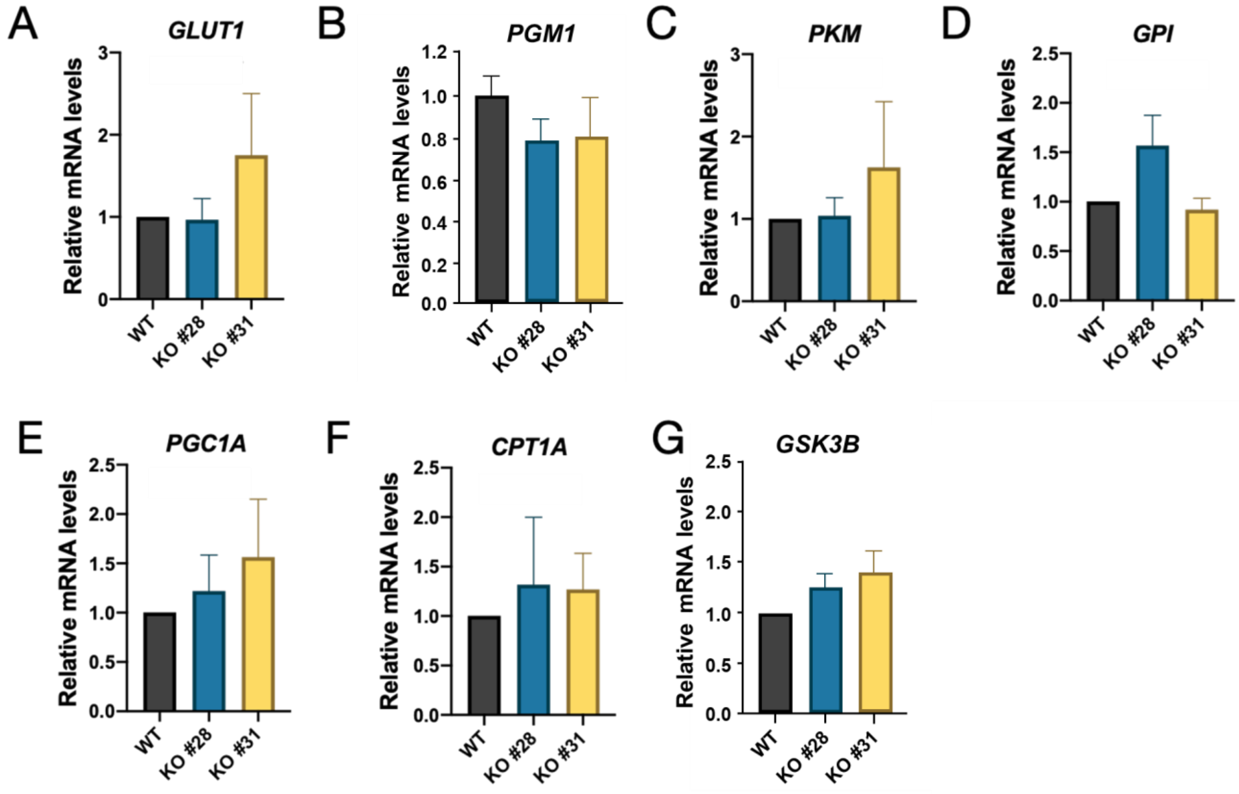

**Figure S2. TRIB2 depletion does not alter the expression of additional glycolytic or mitochondrial metabolic genes.** (A–G) Relative mRNA expression of solute carrier family 2 member 1 (SLC2A1, GLUT1), phosphoglucomutase 1 (PGM1), pyruvate kinase M (PKM), glucose-6-phosphate isomerase (GPI), peroxisome proliferator-activated receptor gamma coactivator 1-alpha (PPARGC1A, PGC1α), carnitine palmitoyltransferase 1A (CPT1A) and glycogen synthase kinase-3 beta (GSK3B), normalized to GAPDH, in parental (WT) and two independent TRIB2-KO clones (#28 and #31) UACC-62 cells (n = 3 independent assays). Values are mean ± SEM of fold change relative to WT. One-way ANOVA was performed and p-values are indicated in each graph.

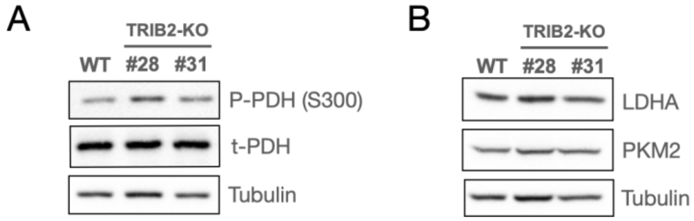


**Figure S3. Depletion of TRIB2 does not alter the levels of proteins involved in metabolism.** Protein levels of phosphorylated (S300) and total PDH (**A**), and LDHA and PKM2 (**B**), in whole cell extracts from parental (WT) and TRIB2-KO (clone #28 and #31) UACC-62 melanoma cell line, using Tubulin as loading control.
